# Antitumor activity of intraperitoneal paclitaxel in orthotopic patient-derived xenograft models of mucinous appendiceal adenocarcinoma

**DOI:** 10.1101/2023.02.01.526672

**Authors:** Ichiaki Ito, Abdelrahman MG Yousef, Princess N Dickson, Zahra A Naini, Michael G White, Karianne G Fleten, Kjersti Flatmark, Keith F Fournier, Natalie W Fowlkes, John Paul Shen

## Abstract

Appendiceal adenocarcinomas (AAs) are a rare and heterogeneous mix of tumors for which few preclinical models exist. The rarity of AA has made performing prospective clinical trials difficult, and in part because of this AA remains an orphan disease with no chemotherapeutic agents approved by the FDA for its treatment. AA has a unique biology in which it frequently forms diffuse peritoneal metastases, but almost never spreads via a hematogenous route and rarely spreads to lymphatics. Given its localization to the peritoneal space we hypothesized that intraperitoneal (IP) delivery of chemotherapy could be an effective treatment strategy. Here we tested the efficacy paclitaxel given by IP administration using three orthotopic PDX models of AA established in NSG mice. Weekly treatment of 25.0 mg/kg of IP paclitaxel dramatically reduced AA tumor growth in TM00351 (81.9% reduction vs. control), PMP-2 (98.3% reduction vs. control), and PMCA-3 (71.4% reduction vs. control) PDX models. Comparing the safety and efficacy of intravenous (IV) to IP administration in PMCA-3, neither 6.25 nor 12.5 mg/kg of IV paclitaxel significantly reduced tumor growth. These results suggest that IP administration of paclitaxel is favorable to IV administration. Given the established safety record of IP paclitaxel in gastric and ovarian cancers, and lack of effective chemotherapeutics for AA, these data showing the activity of IP paclitaxel in orthotopic PDX models of mucinous AA support the evaluation of IP paclitaxel in a prospective clinical trial.

## Introduction

Appendix adenocarcinoma (AA) is a rare disease, diagnosed in approximately 4,000-5,000 Americans each year, but the incidence of AA has dramatically increased in the last ten years, particularly in patients under age 50 (1). With the exception of cases where an appendiceal tumor causes obstruction of the appendix leading to acute appendicitis, AA generally presents only after the tumor has spread to the peritoneal space causing symptomatic peritoneal carcinomatosis. Unique from most other gastrointestinal malignancies, hematogenous and/or lymphangitic metastatic spread of AA is quite rare, meaning that for most patients disease is limited to the peritoneal cavity (2). Complete cytoreductive surgery (CRS) followed by Heated Intraperitoneal Chemotherapy (HIPEC) with either mitomycin C or a platinum agent is currently the best standard-of-care treatment for AA patients; however, many patients, particularly those with high-grade tumors, have tumor involvement in the peritoneal space that is so extensive that they are not candidates for CRS/HIPEC. Given lack of other options, these patients who are not surgical candidates are generally treated with systemic chemotherapy designed for colorectal cancer (CRC), consistent with current National Comprehensive Cancer Network (NCCN) guidelines (3,4). Recent reports have identified clear molecular differences between AA and CRC, and also identified that CRC chemotherapy has limited efficacy in AA relative to CRC (5). Therefore, there is an urgent need to identify novel treatments for AA patients.

Obtaining cytotoxic concentrations of chemotherapy within the peritoneal cavity may be difficult, which may contribute to the limited efficacy of chemotherapy previously observed in AA. Therefore, a regional technique for the treatment of peritoneal carcinomatosis from AA is appealing as theoretically this approach could deliver high concentrations of chemotherapy directly to the tumor while limiting systemic toxicity. There is already a long history of regional peritoneal chemotherapy in the form of HIPEC for the treatment of AA (6) and a more recent trend of using serial pressurized intraperitoneal aerosol chemotherapy (PIPAC) (7,8). The chemical structure of paclitaxel is highly hydrophobic, therefore after IP delivery the drug is slowly absorbed into systemic circulation leading to higher drug exposure in the peritoneal cavity (9). IP administration of taxanes has been considered a promising treatment for eliminating peritoneal metastasis of gastric cancer and has also been used extensively in ovarian cancer (10).

Although to our knowledge serial dosing of intraperitoneal (IP) paclitaxel has never before been performed in patients with appendiceal cancer, for over a decade clinical investigators from Japan have investigated IP paclitaxel as a treatment gastric cancer metastatic to the peritoneum (11–13). Based on a large body of preclinical research, and early phase clinical trials, the Japanese Intraperitoneal Chemotherapy Study Group recently completed the randomized Phase III PHOENIX-GC Trial reporting a 3-year OS rate of 22% in the IP paclitaxel arm (IP paclitaxel plus IV paclitaxel plus PO S-1) versus 6% in the standard chemotherapy arm (IV cisplatin plus PO S-1) (14). A Phase III trial in high grade serous ovarian cancer also demonstrated significant efficacy of IV paclitaxel plus IP cisplatin and IP paclitaxel (IP-therapy), compared to IV paclitaxel plus IV cisplatin (IV-therapy); progression-free survival was 65.6 months in the IP-therapy vs 49.7 months in the IV-therapy, *p* = 0.05 by the log-rank test (15). The efficacy and safety IP paclitaxel in these trials led us to hypothesize that IP paclitaxel would also be active in appendiceal cancer.

## Materials and Methods

### Mice

Female NSG mice, 5-7 weeks of age, were purchased from The Jackson Laboratory (Bar Harbor, ME, USA), and used for tumor implantation. The animals were maintained under specific pathogen-free conditions. Housing and all procedures involving animals were performed according to protocols approved by the Institutional Animal Care and Use Committee of MD Anderson Cancer Center and conducted in accordance with U.S. Common Rule.

### Establishment of PDX models

AA tumor from a flank PDX model was obtained from Jackson laboratory (TM00351). The tumor was cut into ~ 25 - 30 mm^3^ fragments, six of which were placed in the peritoneal cavity, both upper and lower abdominal quadrants, and both flank sides, of NOD/SCID/IL2Rγnull (NSG) mice. The well-being of the mice was carefully monitored, and mice were sacrificed when signs of disease, mainly abdominal distention together with rough coat and reduced mobility, were seen. The tumors in peritoneum were harvested and transferred to subsequent NSG mice. An orthotopic TM00351 PDX model was established after 3 subsequent IP passages. The IP tumor was visualized by a 4.7T small animal MRI system (Bruker Biospin MRI, Billerica, MA, USA), 40 sections of 0.75 mm thickness and 0.25 mm gap of each section, throughout almost entire legion of peritoneal cavity in each mouse. The tumor growth was evaluated by modified peritoneal Response Evaluation Criteria In Solid Tumors (mpRECIST) method, a quantitative measuring system designed for mucinous peritoneal disease, which is the sum of the longest diameter of up to 5 target lesions in the abdominal cavity (NCT01946854) (5,16).

PDX models of pseudomyxoma peritonei from AA, PMP-2 and PMCA-3, were generously provided by Dr. Kjersti Flatmark (University of Oslo), and originally established in the peritoneum of nude mice (17,18). In these PDX models tumor did not spread beyond the peritoneal space. Ascites was obtained from PMP-2 and PMCA-3 using a 3 mL syringe and transferred into NSG mice by IP injection with the syringe attached with 21-gauge needle (100 μl/mouse). The tumors were passaged 3 times to establish PMP-2 and PMCA-3 models in NSG mice. Similar to the tumors in nude mice, the tumor grew as a mucinous ascites in PMP-2 and PMCA-3 in NSG mice, which were diffusely disseminated within the entire peritoneal cavity. Quantitative changes in mucinous peritoneal disease were visualized by MRI. Tumor volume in each mouse was measured by taking the sum of the total area tumor from 5 fixed MRI sections in the peritoneum using OsiriX Lite software v12.0.1 (Pixmeo, Switzerland).

### Paclitaxel treatment

Paclitaxel, dissolved in castor oil, citric acid, and dehydrated alcohol USP, was obtained from residual unused doses from the Mays clinic pharmacy at the University of Texas MD Anderson Cancer Center. Paclitaxel was freshly diluted 5-fold with physiological saline. PDX mice were IP or IV injected with diluted paclitaxel (0.5 ml) with a dose range from 6.25 to 25.0 mg/kg mouse body weight for IP or 6.25 and 12.5 mg/kg for IV. PDX models were treated with paclitaxel weekly (3 weekly treatments and 1 week off, for two cycles) or biweekly (up to 6 biweekly treatments). In a preliminary study, we confirmed that the inactive components in the solvents [castor oil (105.4 mg/ml), citric acid (0.4 mg/ml), and dehydrated alcohol USP (9.94% v/v)] did not have a significant effect on tumor growth or body weight in PMCA-3 PDX mice (data not shown).

### Organoids

TM00351 tumor were dissociated in a tumor digestion media [Advanced DMEM/F12 containing 10 mM HEPES (Gibco, Grand Island, NY, USA), 1x GlutaMax (Gibco), 1x primocin (InvivoGen, San Diego, CA USA), 5 mg/ml collagenase II (Thermo Fisher Scientific, San Jose, CA, USA), 1 mg/ml dispase II (Thermo Fisher Scientific), and 2.5% fetal bovine serum (Hyclone, Logan, Utah, USA)] using gentleMACS Dissociator (Miltenyi Biotec, Bergisch Gladbach, Germany). After the completion of 37_h_TDK_1 program, single cells were obtained from the suspension by passing through 70 μm cell strainer, embedded in Matrigel (BD Biosciences, Bedford, MA, USA; 1 × 10^5^ cells/50 μl dome/well in a 24-well plate), and cultured in organoid growth medium (OGM; Advanced DMEM/F12 containing 50% of L-WRN cell conditioned medium, 1x GlutaMAX, 10 mM HEPES, 100 ng/ml recombinant FGF10 (R&D Systems, Minneapolis, MN, USA), 10 nM Gastrin I (Thermo Fisher Scientific), 1.25 mM N-acetylcysteine (Sigma-Aldrich, St. Louis, MO, USA), 10 mM Nicotinamide (Sigma-Aldrich), and 1x B-27 supplement (Thermo Fisher Scientific). For first 3 days, Y27632 ((Thermo Fisher Scientific; 10 μM) was added to OGM. The culture medium was changed every 3 days to maintain an organoid growth.

To test the efficacy of paclitaxel on the organoid growth, established TM00352 organoids were dissociated by TrypLE™ Express Enzyme (Gibco), and single cells were embedded in Matrigel (100 cells/μl, 5 μl dome/well in 96-well plate). Three days after the cultivation in OGM, the culture medium was changed with OGM supplemented with various doses of paclitaxel. Cells/organoids were incubated in IncuCyte Live-cell analysis system for 3 weeks.

### Statistics

Statistical analysis was carried out using GraphPad Prism 9 (GraphPad Software). An unpaired two-tailed Student’s t-test was used to determine significance of differences between two groups. When appropriate, we have estimated variation within each group of data and ensured that it is similar between groups that are being statistically compared. The log-rank test was used to determine statistically significant differences between two Kaplan-Meier survival curves.

## Results and Discussion

### Phenotypic and histological features of AA tumors in PDX models

Three mucinous AA PDX models, TM00351, PMP-2 and PMCA-3, were established in NSG mice. TM00351, originally established from a tumor diagnosed as metastatic appendix carcinoma, is a poorly differentiated (grade 3, also called high-grade) (19) mucinous adenocarcinoma with *KRAS^G12D^* and *TP53^P72R,V274F^* mutations (**Fig. 1A**). In NSG mice TM00351 formed a mix of solid tumor and a gelatinous, mucinous ascites in the peritoneal cavity typically 2 or 3 months after implantation with histological features consistent with high-grade mucinous appendiceal adenocarcinoma (**Fig. 1B**). PMP-2 was collected from a disseminated abdominal mucinous tumor in a female patient with pseudomyxoma peritonei arising from a mucinous adenocarcinoma of the appendix (17). PMP-2 is a high-grade mucinous AA with *KRAS^G12V^* and *GNAS^R201C^* mutations (**Fig. 1A**) (20). In NSG mice it exhibited a similar histological growth pattern to the human tumor with bland epithelium lining large accumulations of extracellular mucin (**Fig. 1B**). PMCA-3, a high-grade mucinous AA with signet ring features and *BRAF^V600E^* and *GNAS^R201C^* mutations (**Fig. 1A**) (20), was collected from a peritoneal metastasis in a male patient with carcinomatosis presumed to come from a primary AA (18). Histologically in both patient and NSG mouse it was characterized by signet ring cells, either as single cells or in small clusters of tumor cells surrounded by mucin (**Fig. 1B**). For all three models, tumor implanted in the peritoneum did not spread outside of the peritoneal space, even after multiple serial passages. Taken as a whole the similarity in histology and natural history between the human tumors and orthotopic PDX models suggests that orthotopic PDX are an appropriate model to study IP treatment of AA.

**Figure 1.**
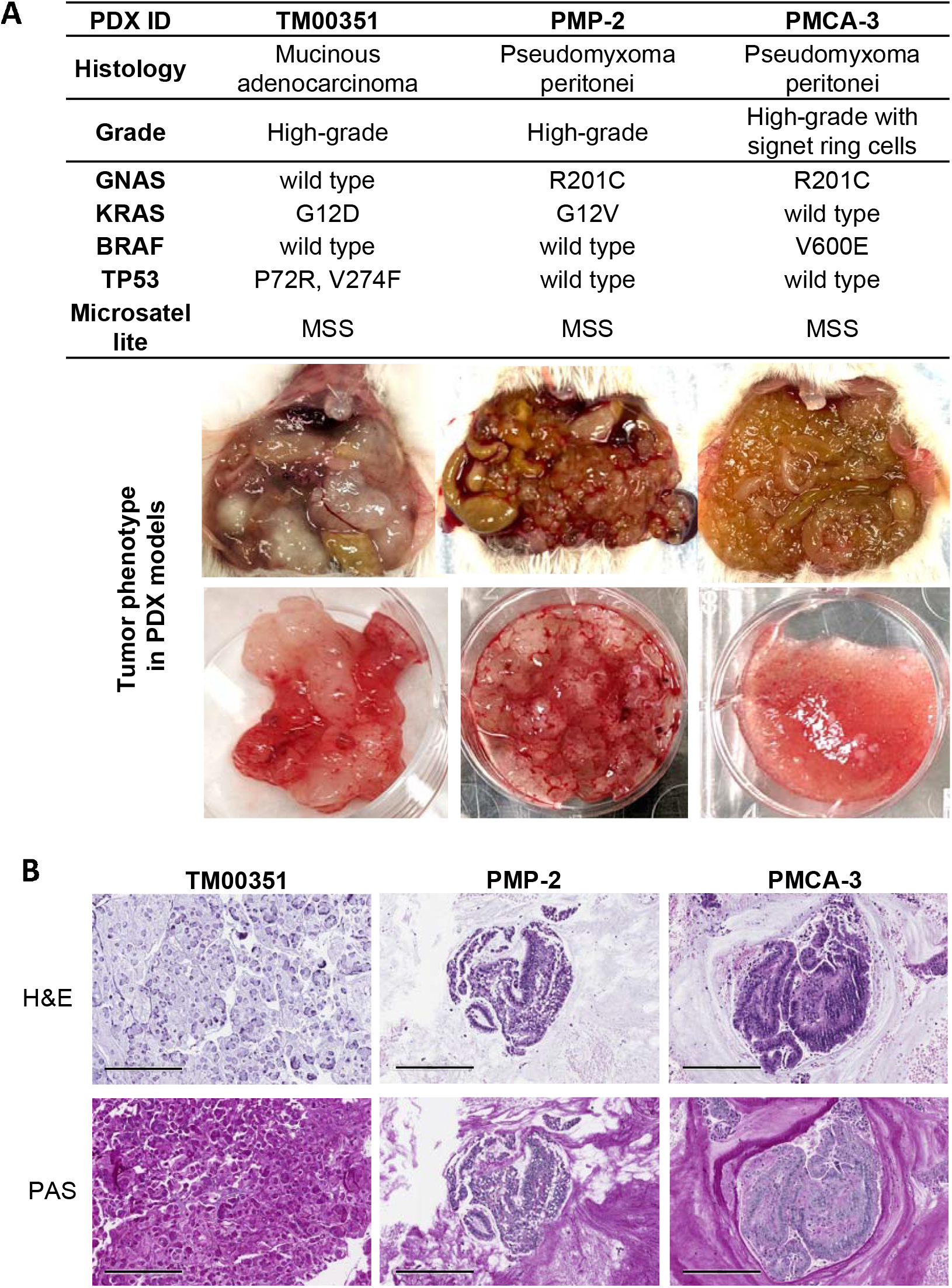
Phenotypic and histological features of AA tumors in PDX models. A) PMP-2 and PMCA-3 models grew as a mucinous ascites, while TM00351 formed a mix of solid tumor and mucinous ascites. PMP-2 and PMCA-3, both with *GNAS^R201C^* mutation, had a mucinous phenotype similar to *GNAS* mutant appendiceal tumors in humans. **B**) Histologically, mucinous adenocarcinoma was seen (Top). Mucinous lakes stain magenta with PAS staining (bottom). Representative images of H&E or PAS-stained slides were taken by Aperio scanner (Leica) and visualized by eSlide Manager (10x). Bar, 300 μm.

### IP paclitaxel suppresses tumor growth in orthotopic models of AA

The efficacy of IP paclitaxel for AA tumor growth in TM00351 PDX model was tested. Tumors uptake was confirmed by MRI 4 weeks after the implantation. Then, mice were treated with 25.0 mg/kg of paclitaxel (human equivalent dose of 70–80 mg/m^2^) (21) via IP injection weekly for 3 weeks followed by one-week rest (**Fig. 2A**). The paclitaxel treatment was repeated one more cycle and tumors were monitored every 4 weeks by MRI. Tumors grew in the peritoneum of control mice injected with IP saline, this growth was greatly impaired in mice treated with IP paclitaxel (**Fig. 2B** & **C**). Tumor growth in the IP paclitaxel group was 81.9% less relative to saline control at 16 weeks (**Fig. 2D**, 3.8 mm in IP paclitaxel vs. 17.0 mm in IP saline); daily tumor growth for IP paclitaxel mice was 88.4% slower than saline group (0.0163 mm/day in IP paclitaxel vs.0.124 mm/day in IP saline). The IP paclitaxel group lost 4.34% of body weight at 1 week after the first treatment (**Fig. 2E**), however, body weight recovered starting from the second week (data not shown). Reduced tumor cellularity and increased necrosis was observed in tumors from IP paclitaxel treated mice (**Fig. 2F**). Interestingly, we observed differential efficacy of IP paclitaxel on TM00351 tumors formed in different loci within the peritoneal cavity (**Supplementary Fig. 1**). The suppressive effect of paclitaxel on tumor growth was also observed *in vitro* in TM00351-derived organoids (**Supplementary Fig. 2**). Taken as a whole these results indicate that IP paclitaxel suppresses the growth the mucinous AA model TM00351.

**Figure 2.**
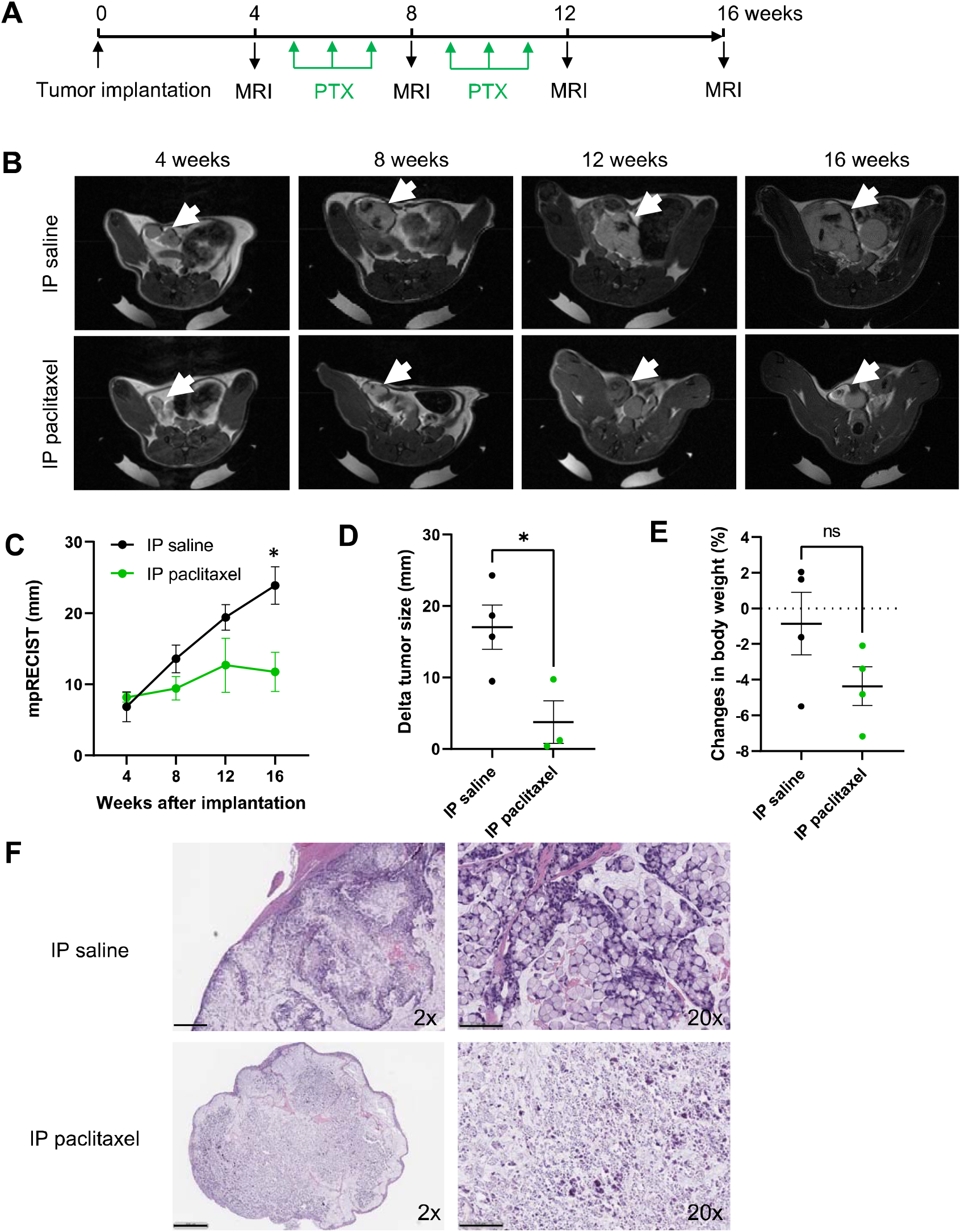
IP paclitaxel suppresses tumor growth of TM00351. **A**) Schedule of the treatment and MRI. Mice were injected IP with paclitaxel (25 mg/kg, weekly for 3 weeks and 1 week rest, and 2 cycles). **B**) MRI of mice abdomen at 4, 8 and 12 weeks after implantation. Arrows indicate tumors in the peritoneal pelvic area. **C-E**) Tumor growth was evaluated by mpRECIST. **C**) Tumor size was scored by mpRECIST. **D**) Tumor growth (mm) from pre-treatment (at 4 weeks) in each mouse was shown. **E**) Changes in body weight compared to pre-treatment are plotted at each time points. **F**) Histological differences in tumors from mice treated with IP paclitaxel and IP saline. Representative images are shown (left, 2x; right 20x). Bars, 500 μm (2x) and 100 μm (20x). *, P < 0.05. ns, not significance.

The efficacy of IP paclitaxel was tested in additional orthotopic models of peritoneal carcinomatosis from mucinous AA. PMP-2 PDX were treated with 25.0 mg/kg of IP paclitaxel weekly for 3 weeks followed by one-week rest, starting from 3 weeks after implantation (**Fig. 3A**). Using mouse MRI to quantitate tumor volumes, as early as two weeks after implantation tumor began to grow in saline control PDX, progressing to diffuse peritoneal carcinomatosis by 6 weeks and requiring mice to be euthanized shortly after week 8. In contrast, for PDX treated with IP paclitaxel, little to no mucinous ascites or tumor was observed either by MRI or direct exploration of the peritoneum of (**Fig. 3B** & **C**). Quantification of tumor volume showed near complete reduction of tumor burden in the IP paclitaxel group (**Fig. 3C**). Mean tumor weight was 4.88 g in saline group vs. 0.09 g in paclitaxel group (98.26% reduction) at 10 weeks (**Fig. 3D**). These results suggest that weekly 25.0 mg/kg IP paclitaxel is highly suppressive for the tumor development of mucinous tumors.

**Figure 3.**
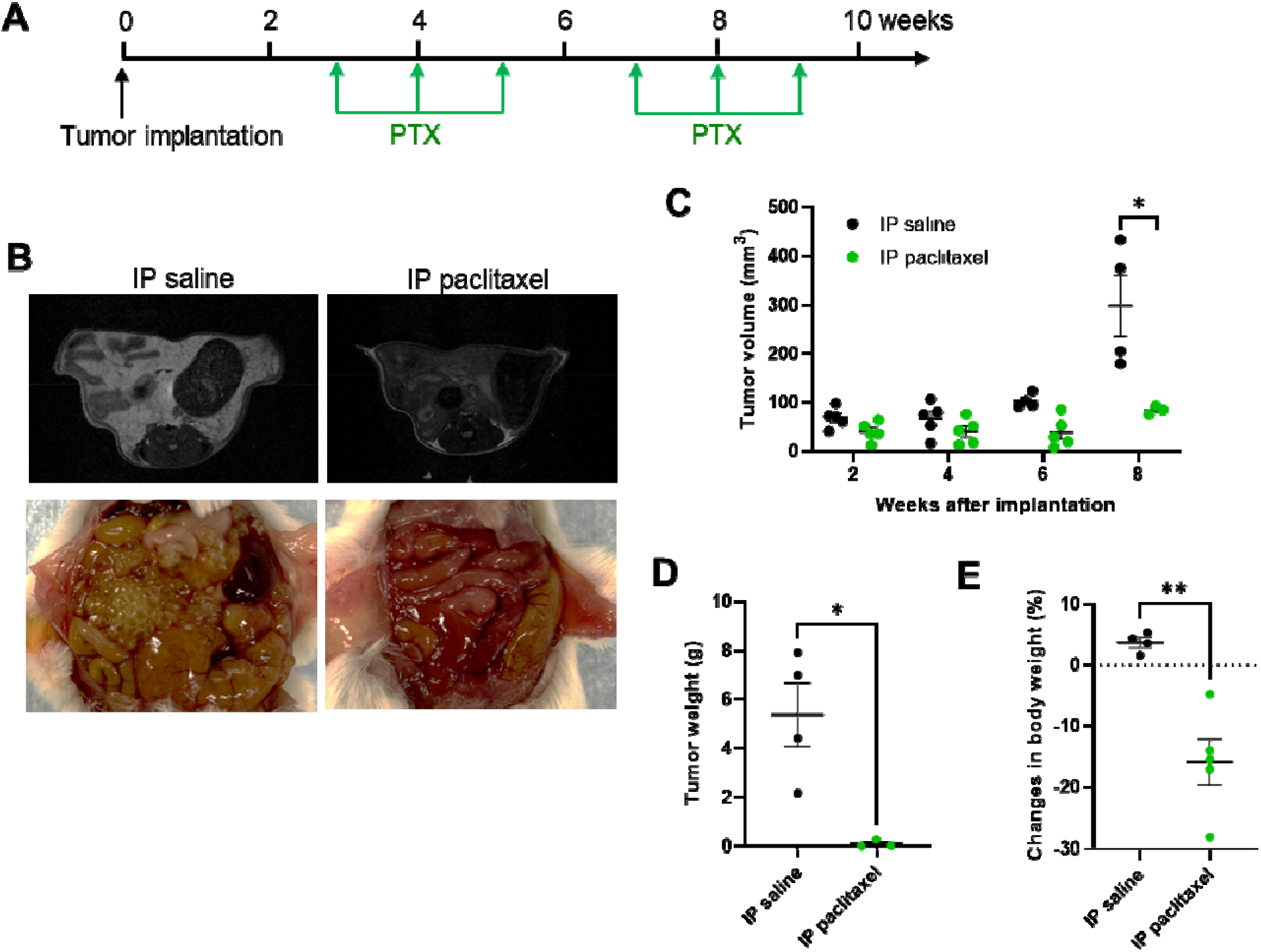
IP paclitaxel treatment in PMP-2 PDX models. **A**) Schedule of the treatment. Mice were injected IP with paclitaxel (25 mg/kg, weekly for **3** weeks and 1 week rest, and 2 cycles). **B**) Representative MRIs at 6 weeks (top) and photos **of** PMP-2 in the peritoneum. **C**) Tumor volume of PMP-2 in the peritoneum. **D**) Tumor weight at **10** weeks. **E**) Percent changes in body weight at each time points are plotted. *, P < 0.05; **, P < 0.01.

### Comparison of IP paclitaxel and IV paclitaxel

To compare the relative efficacy of IP and IV paclitaxel, PMCA-3 PDX mice were treated with 6.25, 12.5, or 25.0 mg/kg of paclitaxel via IP or 6.25 or 12.5 mg/kg via IV, with the same treatment schedule of the experiments with PMP-2 (**Fig. 3A**). PMCA-3 grew as mucinous ascites in the peritoneal cavity and the tumor was readily detected in the peritoneal cavity using MRI (**Fig. 4A**). While 12.5 and 25.0 mg/kg IP paclitaxel were highly effective reducing the tumor burden in the peritoneum [63.2% reduction (*p* = 0.0426) in 12.5 mg/kg IP paclitaxel and 71.4% reduction (*p* = 0.0364) in 25.0 mg/kg IP paclitaxel vs. IP saline at 6 weeks], 6.25 mg/kg IP paclitaxel was moderately effective [20.8% reduction vs. IP saline at 6 weeks, **Fig. 4B**]. Neither 6.25 mg/kg nor 12.5 mg/kg of IV paclitaxel significantly reduced tumor burden in PMCA-3. While 25.0 mg/kg IP paclitaxel reduced body weight by 12.0% one week after the treatment, 12.5 mg/kg IP paclitaxel did not significantly reduce body weight (**Fig. 4C**). Measured at 120 days after implantation (60 days after the last IP paclitaxel treatment), the survival rate of mice treated with 25.0 mg/kg IP paclitaxel was 100% (*p* = 0.0038 and HR = 12.3, vs. IP saline control); for mice treated with 12.5 mg/kg IP paclitaxel survival rate was 33.3% (*p* = 0.0254 and HR = 6.76, vs. IP saline control). All of mice treated with 6.25 mg/kg IP paclitaxel (median survival days of 48 days) and saline control (median survival days of 38 days) were dead within 49 and 77 days, respectively (**Fig. 4D**). None of the tested doses of IV paclitaxel significantly influenced median survival (64 days in 6.25 mg/kg group and 72 days in 12.5 mg/kg group) compared to IV saline control (72 days). H&E staining of tumor sections from PDX treated with 25.0 mg/kg IP paclitaxel revealed reduced cellularity and increased necrosis (**Supplementary Fig. 3**). Given prior reports suggesting that the hydrophobic nature of paclitaxel would cause prolonged retention in the peritoneal space (9), we tested biweekly dosing of IP paclitaxel (25.0 mg/kg) in PMCA-3 PDX mice (**Supplementary Fig. 4A**), average tumor weight in biweekly treated mice was 1.34 g compared to 3.95 g for saline control (66.2% reduction, **Supplementary Fig. 4B**). Of note, biweekly treatment still resulted in body weight loss (14.9%, **Supplementary Fig. 4C**). Biweekly IP paclitaxel treated tumors were histologically similar to those of weekly IP paclitaxel treated tumors (**Supplementary Fig. 4D**). These data indicate that extending IP paclitaxel treatment to biweekly treatment maintains effectiveness.

**Figure 4.**
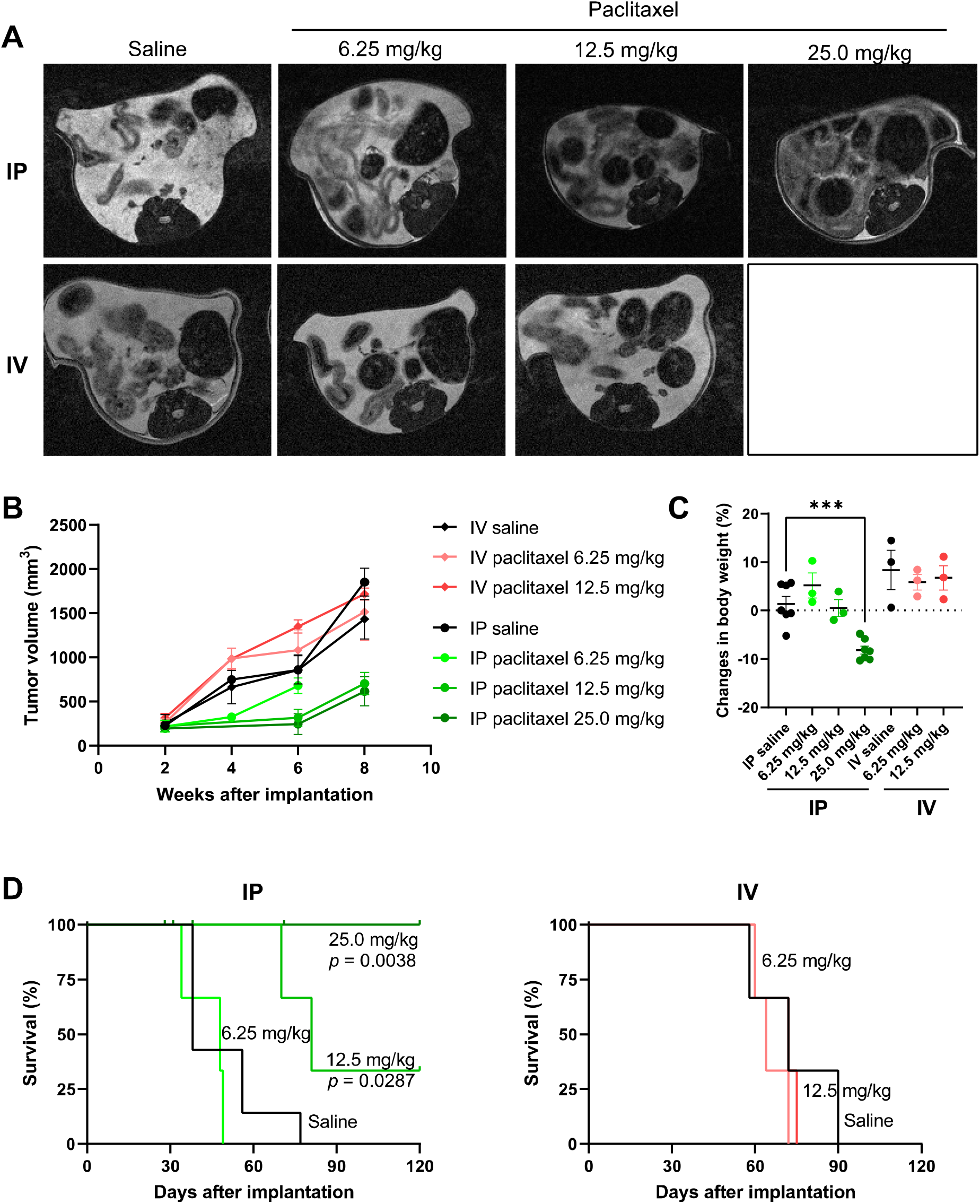
Comparison of IP and IV paclitaxel treatment in PMCA-3 PDX models. Mice were IP or IV injected with paclitaxel (6.25, 12.5 or 25.0 mg/kg, weekly for 3 weeks and 1 week rest, and 2 cycles). **A**) Representative MRIs at 6 weeks (top) are shown. **B**) Quantified tumor volume in the peritoneum. **C**) Percent changes in body weight at each time points are plotted. **D**) Survival of PMCA-3-bearing mice treated with IP or IV paclitaxel. ***, P < 0.001.

In conclusion, we established three orthotopic PDX models of AA, which recapitulate the histology and natural history of appendiceal cancer, and demonstrated that IP paclitaxel is therapeutically active in these models. We find that IP delivery of paclitaxel was more effective relative to IV, with reduced systemic side effects in mice, and that IP paclitaxel treatment extended survival for mice while IV treatment did not. Although less weight loss was observed with IP relative to IV dosing, significant weight loss was seen with higher doses of IP paclitaxel suggesting that further drug modification to further increase the peritoneal concentration would improve therapeutic window. Recently a nanoparticle delivery method using cabazitaxel encapsulated in alginate microspheres has shown promise in this regard (22). Although paclitaxel is active in many gastrointestinal malignancies including gastric, esophageal, and small bowel adenocarcinoma, neither IP nor IV paclitaxel have been prospectively tested in AA, which is likely a legacy of the historic treatment of AA with chemotherapy developed for CRC. *APC* loss-of-function is known to contribute to taxane resistance (23); however, despite being the most commonly mutated gene in CRC, *APC* mutation is uncommon in all subtypes of appendiceal cancer (24). To date, there is not a single drug approved by the Unites States Food and Drug Administration for the treatment of appendiceal cancer. Given these data showing the efficacy of IP paclitaxel in pre-clinical models of appendix cancer and known safety record of IP paclitaxel from multiple prior trials (5) we propose that IP paclitaxel is promising therapeutic strategy that should be tested prospectively in patients with metastatic appendiceal adenocarcinoma.

## Acknowledgments

This work was supported by the National Cancer Institute (L30 CA171000 and K22 CA234406 to J.P.S., and the Cancer Center Support Grant (P30 CA016672), the Cancer Prevention & Research Institute of Texas (RR180035 to J.P.S., J.P.S. is a CPRIT Scholar in Cancer Research), and the Col. Daniel Connelly Memorial Fund. We thank the following core facilities at MD Anderson Cancer Center for their services used in this study: Advanced Technology Genomics Core (Supported by NCI Grant CA016672(ATGC)) and Small Animal Imaging Facility (Supported by the Cancer Center Support Grant CA16672) for *in vivo* live imaging. We also acknowledge Dr. Kenna Shaw and data integration and clinical team members from the Sheikh Khalifa Bin Zayed Al Nahyan Institute for Personalized Cancer Therapy, supported by the Khalifa Bin Zayed Al Nahyan Foundation, for building and maintaining the MOCLIP database.

## Author Contributions

I.I. conceived, designed, performed all experiments and wrote the paper. P.N.D. performed animal experiments, A.M.Y. and Z.A.N. quantified MRI data, K.F.F. and M.G.W. conducted appendiceal tumor preparation, K.G.F. and K.F. provided PMP tumor specimens and counseled the paper writing, and N.W.F. analyzed histological data. J.P.S conceived, designed and supervised the study, and wrote the paper.

## Data Availability

All data supporting the findings of this study are available within the paper and its supplementary information files will be uploaded to GEO.

