## Supplementary Figures for "Antitumor activity of intraperitoneal paclitaxel in orthotopic patient-derived xenograft models of mucinous appendiceal adenocarcinoma"

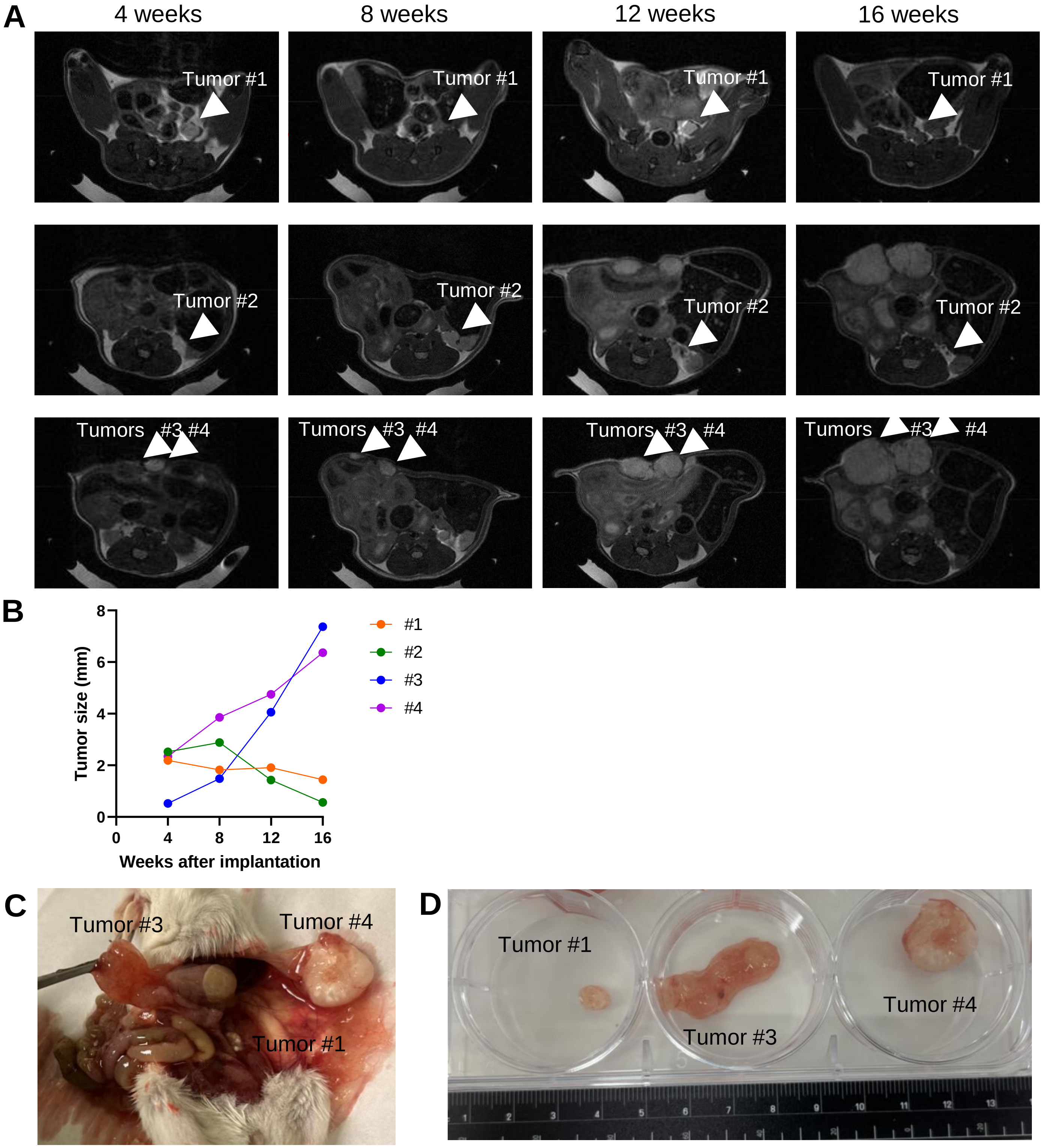


**Supplementary Figure 1. Differential suppressive effect of IP PTX on peritoneal TM00351 tumor growth**

**A**) The mouse was implanted with 6 pieces of tumors in the peritoneum. Tumors #1 and #2 were formed in the pelvic area (#1) and around left ovary region (#2) in the peritoneal cavity. Tumors #3 and 4 were unexpectedly formed on the peritoneal membrane, where the incision were made for the implantation. MRI was taken from mouse treated with weekly IP PTX at a dose of 25 mg/kg 4 to 16 weeks after tumor implantation. See the detailed treatment schedule in Figure 2A. Arrow heads indicate each tumors (tumor #1 to #4). **B**) Tumor size was measured in MRI. **C** and **D**) Tumors were taken from the mouse 180 days after implantation.

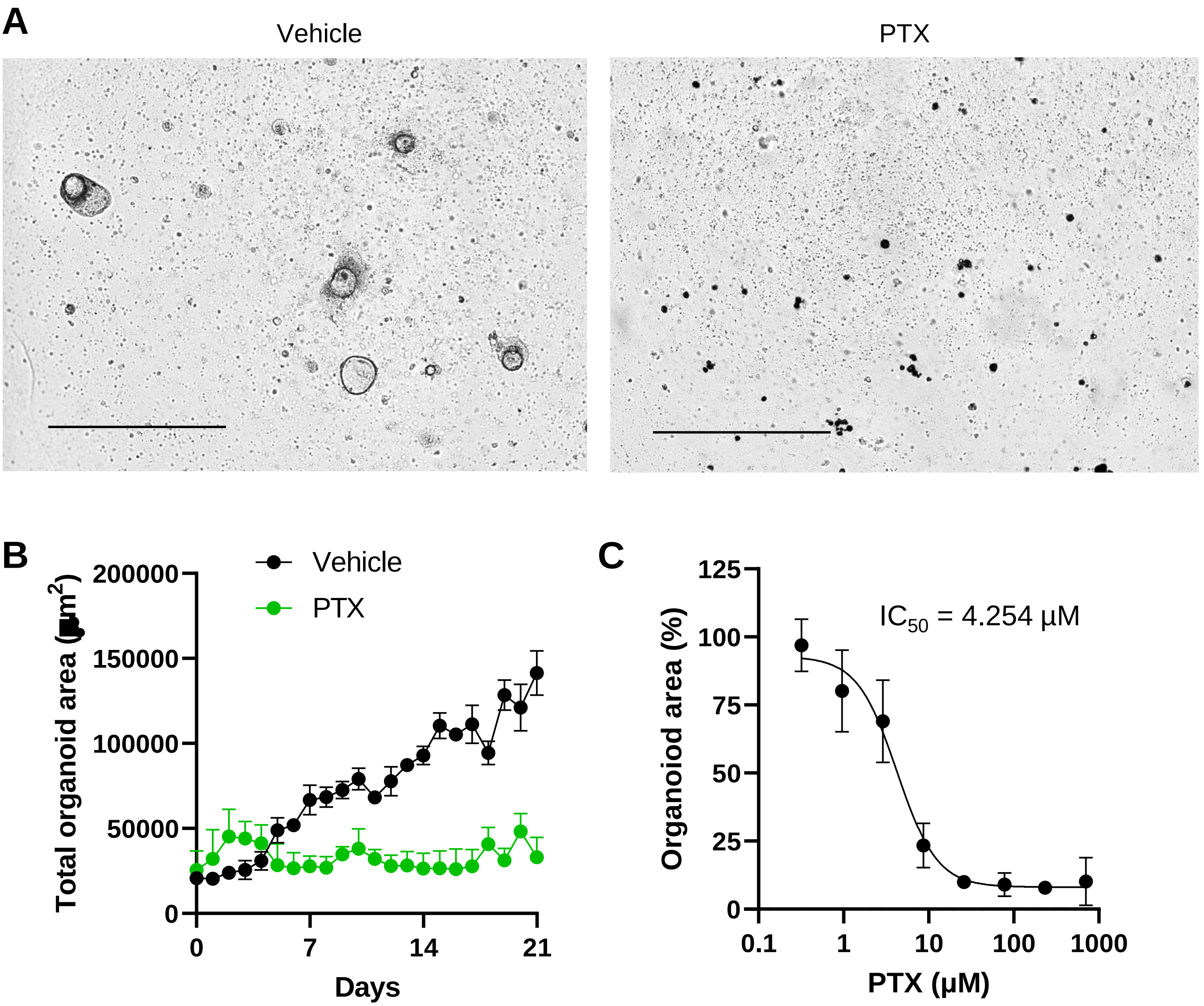


**Supplementary Figure 2. PTX inhibits the growth of organoids derived from TM00351 tumor**
**A**) Tumor cells (1000 cells/wells, 96-well plate) from TM00351 were embedded in Matrigel and cultured in human organoid growth medium (OGM). The organoids were cultured in Incucyte for 21 days. Representative phase images of organoids cultured in OGM added with paclitaxel (8.68 µM) are shown. As a control, the organoids were cultured in OGM with vehicle in the same manner. Bar, 800 µm. **B**) Total organoid area per well were quantified. **C**) Total organoid areas/well after cultivation in OGM with various concentrations of paclitaxel for 21 days were plotted. The half maximal inhibitory concentration ($IC_{50}$) was calculated from relative total organoid area which normalized by that of vehicle control.

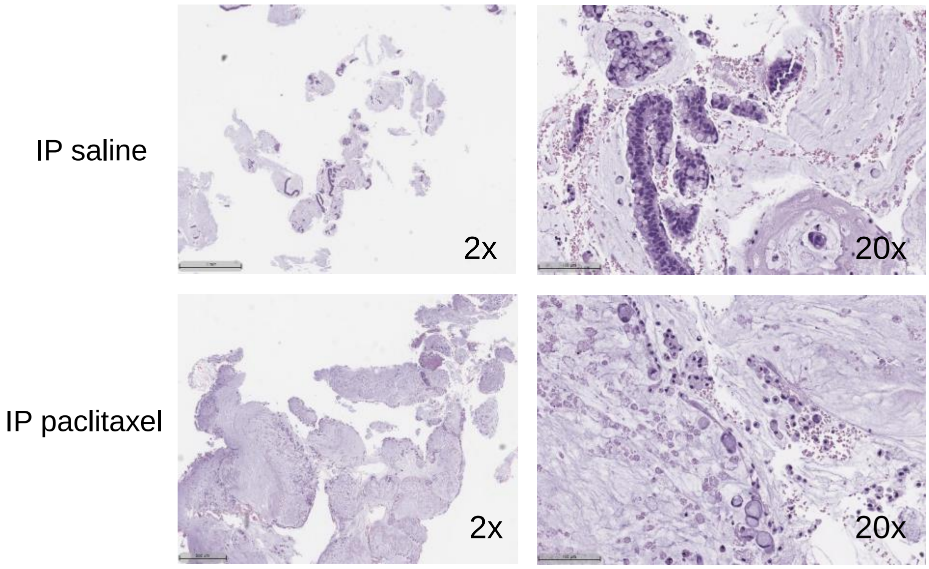


**Supplementary Figure 3. Histology of IP PTX-treated PMCA-3 PDX tumor**

Histological differences in tumors from mice treated with IP paclitaxel and IP saline. Representative images are shown (left, 2x; right 20x).

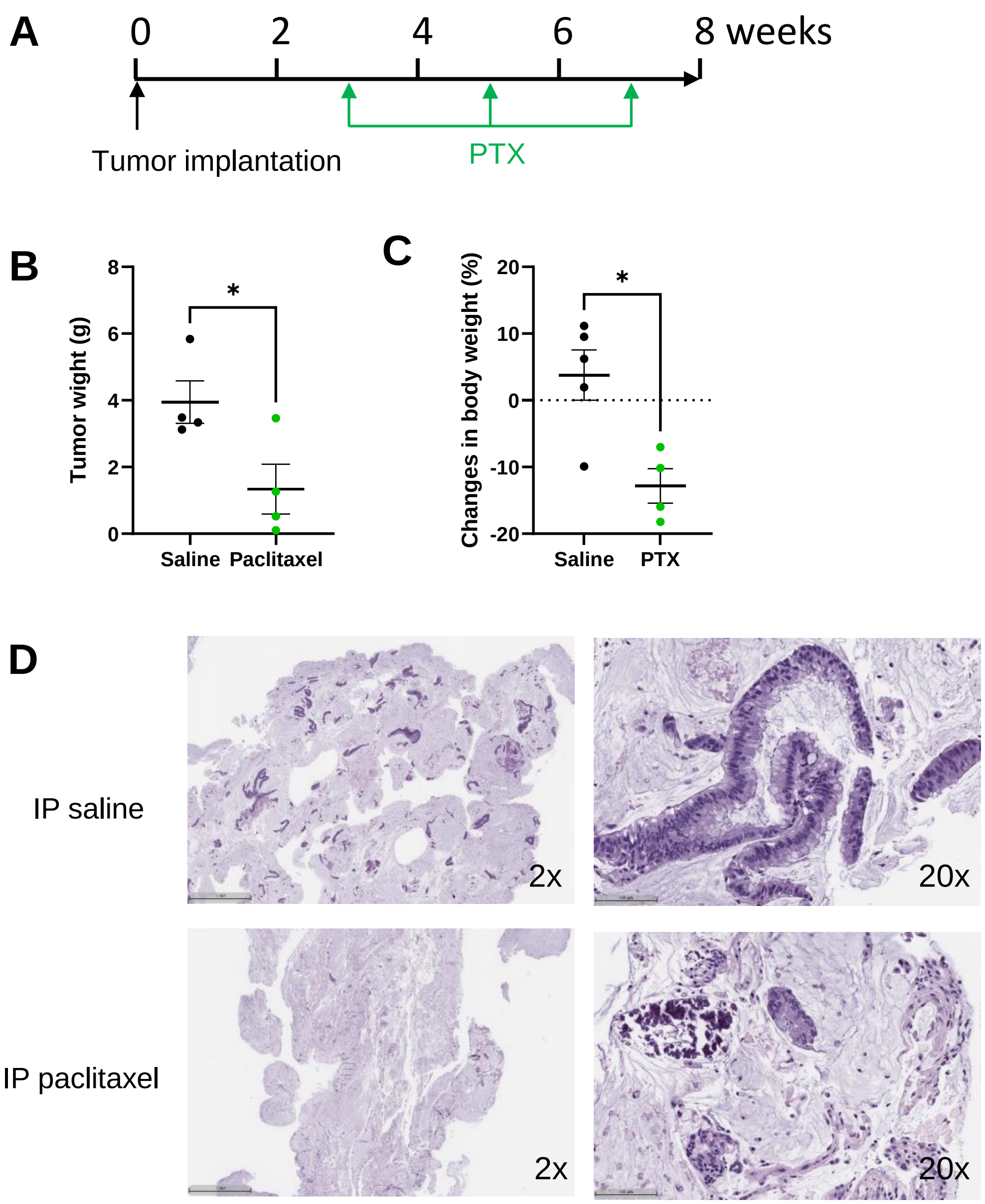


**Supplementary Figure 4. IP PTX biweekly treatment in PMCA-3 PDX model**

**A**) Schedule of the treatment. Mice were IP injected with paclitaxel (25.0 mg/kg, biweekly for 3 weeks). **B**) Tumor weight of PMCA-3 at end point. **C**) Percent changes in body weight at 1 week after IP treatment was plotted. **D**) Histological differences in tumors from mice treated with IP paclitaxel and IP saline. Representative images are shown (left, 2x; right 20x).
